# Lac-Phe mediates the anti-obesity effect of metformin

**DOI:** 10.1101/2023.11.02.565321

**Authors:** Shuke Xiao, Veronica L. Li, Xuchao Lyu, Xudong Chen, Wei Wei, Fahim Abbasi, Joshua W. Knowles, Shuliang Deng, Gaurav Tiwari, Xu Shi, Shuning Zheng, Laurie Farrell, Zsu-Zsu Chen, Kent D. Taylor, Xiuqing Guo, Mark O. Goodarzi, Alexis C. Wood, Yii-Der Ida Chen, Leslie A. Lange, Stephen S. Rich, Jerome I. Rotter, Clary B. Clish, Usman A. Tahir, Robert E. Gerszten, Mark D. Benson, Jonathan Z. Long

**Author notes:** These authors contributed equally (Mark D. Benson, Jonathan Z. Long).

## Abstract

Metformin is a widely prescribed anti-diabetic medicine that also reduces body weight. The mechanisms that mediate metformin’s effects on energy balance remain incompletely defined. Here we show that metformin is a powerful pharmacological inducer of the anorexigenic metabolite Lac-Phe in mice as well as in two independent human cohorts. In cell culture, metformin drives Lac-Phe biosynthesis via inhibition of complex I, increased glycolytic flux, and intracellular lactate mass action. Other biguanides and structurally distinct inhibitors of oxidative phosphorylation also increase Lac-Phe levels *in vitro*. Genetic ablation of CNDP2, the principal biosynthetic enzyme for Lac-Phe, in mice renders animals resistant to metformin’s anorexigenic and anti-obesity effects. Mediation analyses also support a role for Lac-Phe in metformin’s effect on body mass index in humans. These data establish the CNDP2/Lac-Phe pathway as a critical mediator of the effects of metformin on energy balance.

---

Metformin is a widely prescribed anti-diabetic medicine^1,2^. In addition to its glucose lowering activity, many clinical studies have also demonstrated that metformin reduces body weight^3–5^. The effects of metformin on energy balance appears to be due to a suppression of food intake rather than changes in nutrient absorption or energy expenditure^6,7^. Metformin administration to mice also decreases food intake and body weight, demonstrating conservation of its effects on energy balance across mammalian species^8^. The proposed mechanisms by which metformin regulates glucose homeostasis, such as inhibition of endogenous glucose production in the liver, do not appear to account for the effects of this drug on systemic energy balance.

In recent years, focus has turned to the GDF15/GFRAL pathway as a potential mediator of the effects of metformin on body weight^9–12^. GDF15 is a powerful anorexigenic hormone that acts via GFRAL, its cognate hindbrain-restricted receptor, to suppress feeding and obesity in mice^13–17^. Multiple research groups have consistently reported increases in GDF15 levels following metformin treatment in both mice and humans^9–12,18^. Several papers^9,10,12^ also demonstrate that metformin’s effects on body weight are diminished or completely abolished in mouse and rat models with genetically or pharmacologically disrupted GDF15/GFRAL pathways. However, Klein et al.^11^ dispute these results and claim that the GDF15/GFRAL pathway is not required for metformin’s effects on energy balance. As a result, there is ongoing debate about the precise molecular mechanisms that mediate metformin’s anorexigenic and anti-obesity effects.

We recently reported that a blood-borne lactate-derived metabolite called Lac-Phe robustly suppresses food intake and body weight when administered to mice^19^. Lac-Phe is one of the most significantly elevated metabolites in multiple animal models of sprint exercise. Genetic ablation of CNDP2, the principal biosynthetic enzyme for Lac-Phe, reduces basal and exercise-inducible Lac-Phe levels and renders mice hyperphagic and obese after a combination of high fat diet and treadmill running training. These mouse knockout studies provide genetic evidence for the physiologic contribution of the CNDP2/Lac-Phe pathway to energy balance, especially after exercise training. In humans, Lac-Phe is also one of the most inducible metabolites following an acute cardiopulmonary treadmill test. In addition, exercise-inducible Lac-Phe predicts adipose tissue loss during endurance training in overweight and obese individuals^20^.

Beyond sprint exercise, little is known about pharmacological or physiologic stimuli that increase circulating Lac-Phe levels. Here, we identify metformin a strong pharmacological inducer of Lac-Phe levels in sedentary mice and in two independent human cohorts. The effect of metformin to increase Lac-Phe is cell-autonomous and occurs in a CNDP2-dependent manner in macrophages and gut epithelial cells, but not hepatocytes. Mechanistically, metformin inhibits complex I, increases glycolytic flux, and drives Lac-Phe biosynthesis via intracellular lactate mass action. Statistical mediation analyses demonstrate that Lac-Phe contributes to the anti-obesity effects of metformin in humans. Using global CNDP2-KO mice that are deficient in Lac-Phe biosynthesis, we show that induction of Lac-Phe by metformin is critical for mediating its anorexigenic and anti-obesity effects. Interestingly, CNDP2-KO mice remain fully sensitive to metformin’s anti-diabetic effects. CNDP2-KO mice also remain fully sensitive to other anorexigenic agents including agonists that themselves do not stimulate glycolytic flux. These data demonstrate that the CNDP2/Lac-Phe pathway mediates the effects of metformin on energy balance.

## Results

### Effect of metformin treatment on plasma Lac-Phe in humans and mice

To determine if metformin increases Lac-Phe in humans, we used liquid chromatography-mass spectrometry (LC-MS) to measure plasma Lac-Phe levels from individuals who had not previously received metformin therapy and were randomized to receive either metformin or placebo. The original study was conducted from 1998-2004 at Stanford Medicine and consisted of participants with type 2 diabetes (N = 31, mean age 58 ± 10 years; mean BMI, 29.4 ± 4.6 kg/m^2^)^21^. Additional baseline characteristics of this cohort are detailed in Methods section. “Pre” and “post” samples were collected in the fasted state just prior to and 12-weeks following, respectively, initiation of metformin. In this protocol, the starting metformin dose was 500 mg twice daily with a slow up-titration to a final dose of 2 g daily. As expected, we found that levels of metformin were dramatically elevated in all individuals randomized to metformin (Fig. 1A). In addition, we observed robust increases in plasma Lac-Phe levels (mean ± SEM, pre 0.09 ± 0.03 μM, post 0.32 ± 0.11 μM, P = 0.023) (Fig. 1B). No changes were observed in plasma lactate levels with metformin therapy (mean ± SEM, pre 0.26 ± 0.02 μM, post 0.31 ± 0.03 μM, P = 0.12) (Fig. 1C).

**Figure 1.**
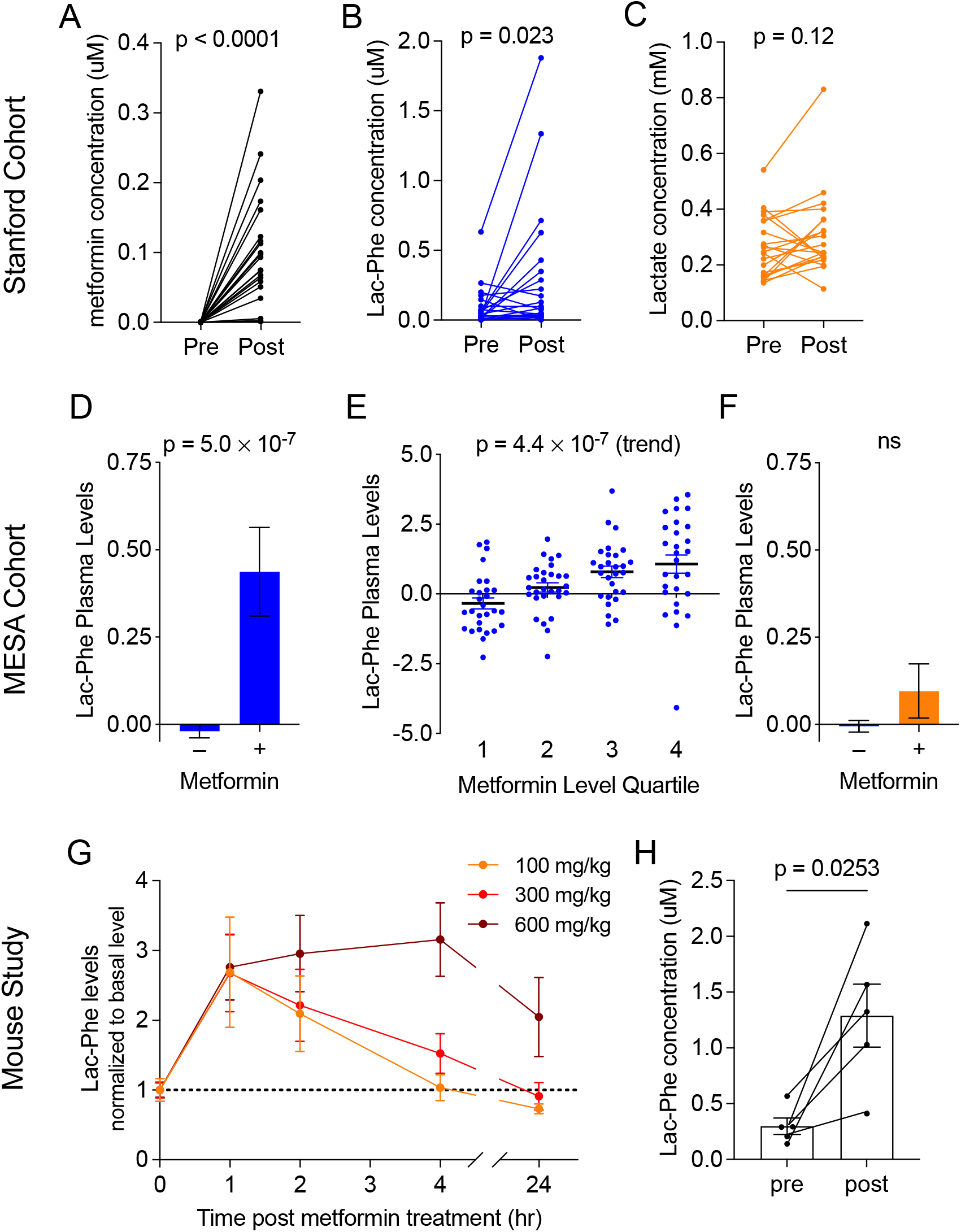
Metformin increases plasma Lac-Phe levels in humans and mice. (A-C) Plasma levels of metformin (A), Lac-Phe (B), and lactate (C) pre- and post-3 months of metformin treatment in the Stanford cohort (n = 21). (D) Participants in MESA on metformin (n = 179) had higher plasma levels of Lac-Phe compared to participants not on metformin (n = 3477). (E) Among the 179 participants on metformin, a significant stepwise increase in plasma levels of Lac-Phe was detected with increasing quartiles of metformin. (F) No significant difference in plasma levels of lactate in participants on metformin compared to participants not on metformin. (G) Circulating Lac-Phe levels at 0, 1, 2, 4, 24 hrs post metformin oral gavage (n = 5). (H) Circulating Lac-Phe concentrations pre- and 1 hour post-300 mg/kg metformin oral gavage (n = 5). P values in (A-C), and (H) were calculated with paired t tests. P values in (D) and (F) were generated using linear regression models adjusting for age, sex, fasting plasma glucose, total cholesterol, and hypertension status. P value in (E) was generated using the Jonckheere-Terpstra test for trend. Data are shown as mean ± SEM.

To further investigate the association of metformin with Lac-Phe levels in a larger population-based human cohort, we next analyzed the association of metformin use with plasma Lac-Phe levels in participants of the Multi-Ethnic Study of Atherosclerosis (MESA)^22^. We recently measured circulating levels of Lac-Phe, lactate, and metformin in plasma samples from N=3,656 participants of MESA using LC-MS. The baseline characteristics of the MESA study participants are detailed in Table S1, and included 179 participants on metformin. In multivariable regression analyses (adjusted for age, sex, fasting glucose, total cholesterol, and hypertension status), participants on metformin had significantly higher circulating levels of Lac-Phe compared to participants who were not on metformin (β = 0.53, P = 5.0×10^−7^) (Fig. 1D). The measurement of metformin levels in parallel to Lac-Phe levels by LC-MS allowed us to directly relate plasma levels of these two metabolites in a human population. Among the 179 MESA participants on metformin, we observed a significant step-wise increase in circulating Lac-Phe levels across increasing quartiles of plasma metformin levels (z = 5.05, P(trend) = 4.4×10^−7^) (Fig 1E). By contrast, we did not observe significantly higher levels of lactate in participants on metformin compared to participants not on metformin (β = 0.08, P = 0.47) (Fig. 1F).

To investigate the effects of oral metformin administration on Lac-Phe in mice, we administered metformin (100, 300, or 600 mg/kg, PO) to wild-type C57BL/6J diet-induced obese (DIO, 14-15 weeks old) mice and performed LC-MS on blood plasma. Plasma metformin levels achieved ∼2-10 μM over a period of at least four hours, which returned back to baseline by 24 h (Fig. S1A). In these same samples, we observed a robust induction of Lac-Phe levels that peaked at 1 h post-administration and persisted for up to 4-24 hrs, depending on the dose (Fig. 1G). Absolute quantitation of Lac-Phe showed baseline levels of 0.30 ± 0.07 μM (mean ± SEM) and metformin-inducible levels of 1.29 ± 0.28 μM (mean ± SEM) at the 300 mg/kg dose (Fig. 1H). The increase in Lac-Phe after a single administration of metformin is comparable in magnitude, and much greater in duration, to what was previously observed after a single bout of sprint exercise in mice^19^. We conclude that metformin administration increases plasma Lac-Phe levels in both mice and humans.

### Metformin increases extracellular Lac-Phe in vitro in a CNDP2-dependent manner

To investigate the cellular source of elevated Lac-Phe after metformin treatment, we measured Lac-Phe levels in conditioned media after metformin treatment across a large panel of cell lines and primary cells. Among this set of cells were primary mouse hepatocytes as well as human and mouse hepatocyte cell lines (HepG2 and AML12), as well as other, CNDP2+ cells that we had previously shown to exhibit robust Lac-Phe production and secretion (e.g., primary macrophages). Cells were treated with increasing concentrations of metformin overnight and Lac-Phe was measured by LC-MS. The strongest induction of Lac-Phe production by metformin was seen in primary macrophages, followed by Caco-2, a human gut epithelial cell line and BV-2, a mouse microglial cell line (Fig. 2A). By contrast, in the other 8 cell lines examined, little metformin-stimulated Lac-Phe production was observed (Fig. 2A).

**Figure 2.**
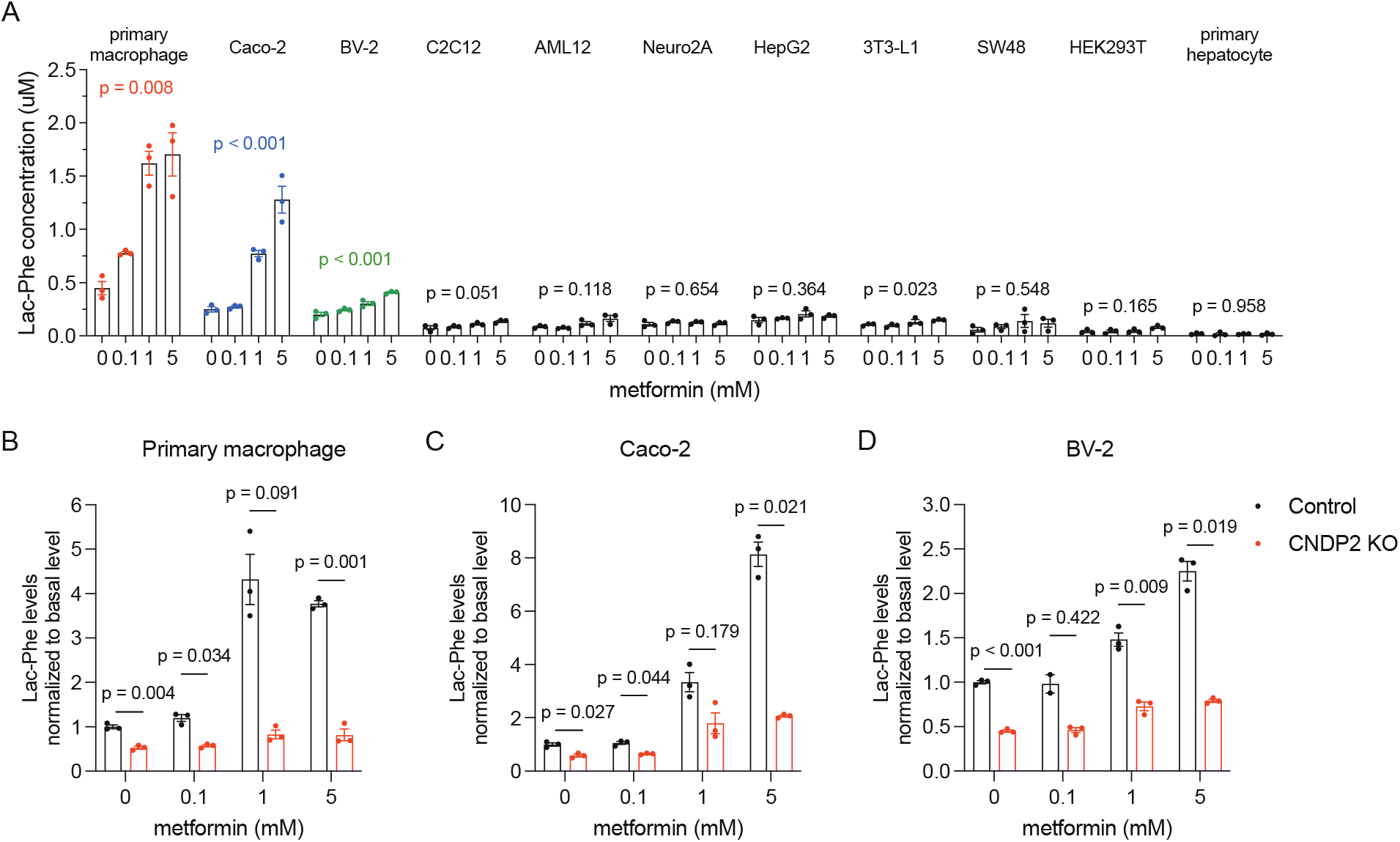
Metformin induced CNDP2-dependent Lac-Phe production *in vitro*. (A) Lac-Phe concentrations in conditioned media from various cell lines and primary cells when treated with metformin at doses indicated below. (B-D) Lac-Phe concentrations compared to basal Lac-Phe production in wild-type primary macrophages (B), Caco-2 cells (C), and BV-2 cells (D) with metformin treatment. N = 3 for all groups. P values were calculated with Welch’s ANOVA test in (A) and multiple unpaired t tests with Welch and Bonferroni-Dunn corrections in (B-D).

We next tested if such metformin stimulated Lac-Phe production also genetically depends on CNDP2. Primary mouse thioglycolate-elicited peritoneal macrophages were directly isolated from CNDP2-KO mice. Ablation of CNDP2 in Caco-2 and BV-2 cell lines was achieved via the CRISPR-Cas9 technique. Knockout of CNDP2 in all three cells was verified by loss of immunoreactivity by Western Blot using an anti-CNDP2 antibody (Fig. S2). In CNDP2-KO primary macrophages, Lac-Phe production was reduced by ∼50% in basal state, and further reduced by ∼80% in metformin stimulated conditions (Fig. 2B). A similarly large diminution of metformin-stimulated Lac-Phe production was also observed in CNDP2-KO Caco-2 and BV-2 cell lines (Fig. 2C,D). We conclude that metformin cell autonomously increases Lac-Phe production in a CNDP2-dependent manner in multiple cell types, including macrophages and gut epithelial cells, but not in hepatocytes.

### Metformin drives Lac-Phe biosynthesis via complex I inhibition and lactate mass action

Metformin is direct inhibitor of mitochondrial complex I. Such a mechanism and molecular target might explain metformin’s induction of Lac-Phe due to the subsequent glycolytic shift and increased lactate flux from intracellular lactate.^19,23^ To directly test this prediction, we first measured cellular respiration using a Seahorse respirometer in primary macrophages and Caco-2 cells treated with increasing doses of metformin. As expected^24,25^, metformin dose-dependently inhibited cellular respiration (Figs. 3A, S3A). Importantly, the concentrations at which metformin inhibited respiration coincided with that at which metformin increased Lac-Phe levels (∼ 1 mM), providing correlational evidence that metformin engagement and inhibition of complex I is associated with its ability to increase Lac-Phe levels in cells.

**Figure 3.**
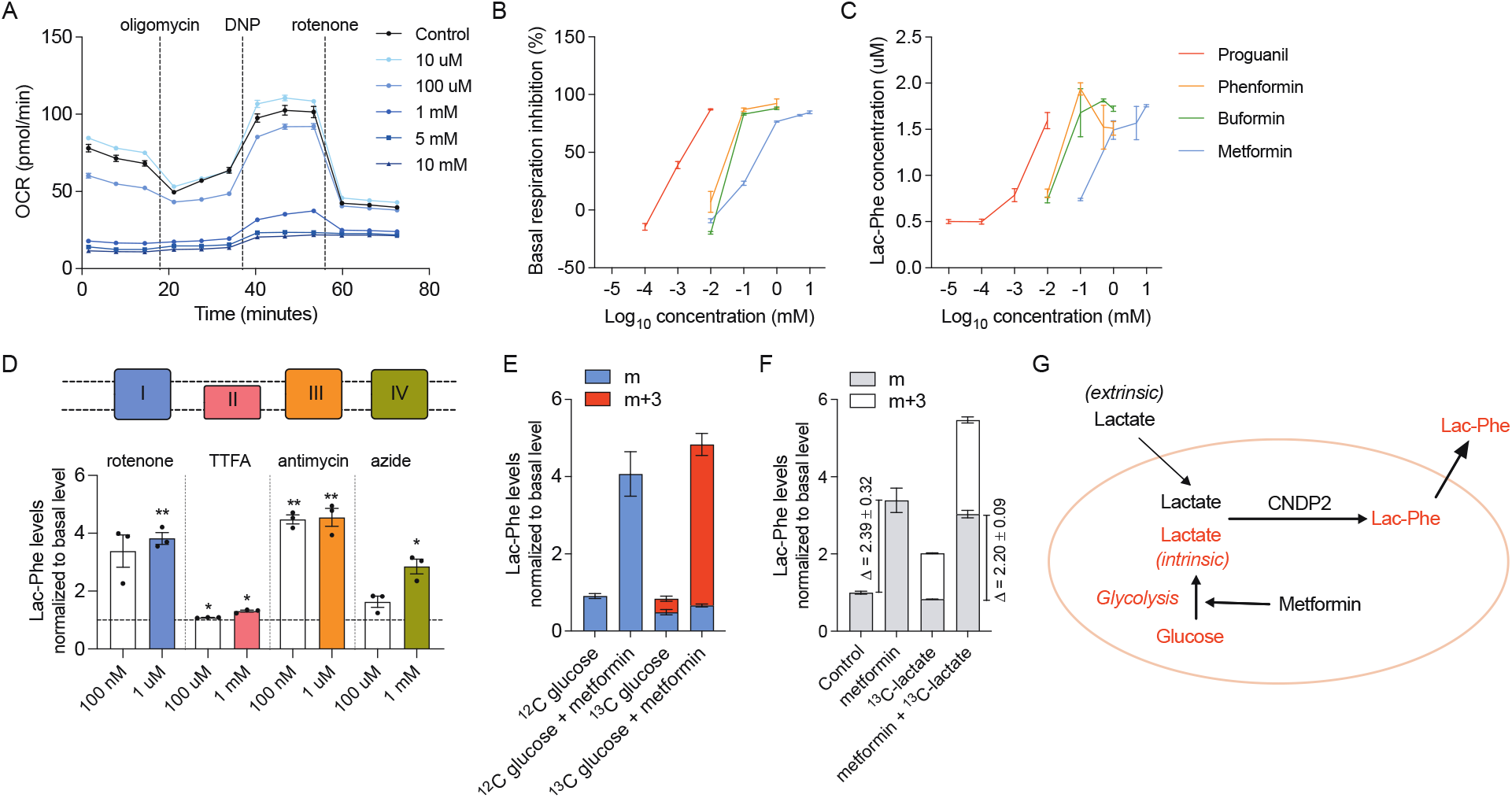
Increase of Lac-Phe production by increased glycolytic flux. (A) Metformin reduces basal and maximal mitochondrial respiration in primary macrophages (n = 5). (B) Biguanides inhibits basal mitochondrial respiration in primary macrophages (n = 5). (C) Biguanides increase Lac-Phe production in primary macrophages (n = 3). (D) Mitochondrial complex inhibitors increase Lac-Phe production in primary macrophages (n =3). (E) Metformin increases ^13^C-labeled Lac-Phe production from ^13^C-labeled glucose in primary macrophages (n = 3). (F) Metformin increases Lac-Phe production mainly through intracellular lactate mass action (n = 3). (G) Schematic illustration of metformin-induced Lac-Phe production. P values in (D) were calculated using one sample t test. * p < 0.05, ** p < 0.01.

Other structurally related biguanides, such as buformin, phenformin, and proguanil, are also complex I inhibitors, but with different potencies compared to metformin. We therefore tested whether these other biguanides increased extracellular Lac-Phe levels in primary macrophages, and if so, whether the concentrations at which the biochemical effect was observed also coincided with the concentrations needed to suppress cellular respiration. As shown in Fig. 3B and Fig. S3B, buformin, phenformin, and proguanil suppressed cellular respiration at concentrations of 1 μM, 1 mM, and 1 mM respectively in primary macrophages; we observed a corresponding increase in extracellular Lac-Phe at these same concentrations (Fig. 3C). These data show that the concentrations of biguanides needed for their inhibition of complex I correlates with their ability to induce Lac-Phe levels, and provides additional evidence that metformin and other biguanides increase Lac-Phe via inhibition of complex I.

Our model for metformin induction of Lac-Phe would also predict that any pharmacological inhibition of oxidative phosphorylation should also increase Lac-Phe levels by inducing a glycolytic shift to drive intracellular lactate mass action. We therefore treated primary macrophages with rotenone, TTFA, antimycin, and azide, which are structurally and mechanistically inhibitors of mitochondrial oxidative phosphorylation (inhibition of complex I, II, III, and IV, respectively). Across all of these treatments, we observed 1.5-5-fold increases in extracellular Lac-Phe levels. Therefore pharmacological suppression of oxidative metabolism is a general mechanism that induces Lac-Phe levels in cells.

We previously showed that in the context of sprint exercise, Lac-Phe biosynthesis is driven by mass action via increases in extracellular, muscle-derived lactate. The absence of a robust increase in circulating lactate after metformin treatment suggests that a potentially distinct source of lactate drives Lac-Phe biosynthesis under these conditions. We therefore tested whether intracellular (rather than extracellular) lactate is a major contributor to Lac-Phe biosynthesis after metformin treatment. To directly determine such a contribution, we measured ^13^C-labeled glucose incorporation into Lac-Phe in the basal state and after treatment with metformin. As shown in Fig. 3E, the majority (>80%) of the metformin-inducible increase in Lac-Phe was of the m+3 isotope, which is expected given that intracellular lactate derived from glycolysis of ^13^C-labeled glucose would have three heavy carbon labels. We also tested the effect of metformin on Lac-Phe production in the presence of extracellular ^13^C-labeled lactate. While metformin once again increased total extracellular Lac-Phe levels, there was no substantial change in the m+3 isotope fraction, demonstrating minimal contribution of extracellular lactate to metformin-inducible Lac-Phe production (Fig. 3F). These data directly demonstrate that metformin increases the flux of intracellular, but not extracellular lactate to drive Lac-Phe production. We therefore conclude that whereas sprint exercise drives Lac-Phe production via extracellular lactate mass action, metformin drives Lac-Phe production via intracellular lactate mass action via glycolysis-derived lactate (Fig. 3G).

### Contribution of Lac-Phe to metformin’s effects on BMI in humans

Because of the previously reported effects of Lac-Phe on food intake, we used statistical mediation methods to investigate if metformin-inducible Lac-Phe levels might mediate, at least in part, the anti-obesity effects of metformin in participants of MESA. Our measurement of baseline Lac-Phe plasma levels in participants of MESA allowed for these analyses since longitudinal measurements of body mass index (BMI) over time were available in this cohort. Specifically, we sought to examine the contribution of Lac-Phe to the relationship between baseline metformin use (exposure) and change in BMI (outcome) over the study period (mean follow-up of 9.46 years). A straightforward application of mediation methods to the entire cohort, however, proved infeasible because all participants on average gained a modest amount of weight during the study period (ΔBMI 0.24 ± 0.04 kg/m^2^, N = 3645). We therefore performed a post-hoc subgroup analysis on two populations of MESA participants: those participants who gained weight (ΔBMI 1.84 ± 0.04 kg/m^2^, N = 2026) and those participants who lost weight (ΔBMI −1.78 ± 0.05 kg/m^2^, N = 1619) over the study period. Baseline characteristics of participants in each subgroup are detailed in Table S2.

Among MESA participants who lost weight during the study period and who had available metformin, Lac-Phe, and lactate levels measured by LC-MS (N = 1181), we observed a significant association between metformin use and BMI using age- and sex-adjusted linear regression models (β = −0.52, P = 4.8×10^−2^). This association formed the basis for the total effect between metformin use and BMI in an “unmediated model” (Fig. 4A). We next constructed a “mediated model” that connected metformin use to BMI through the effect of Lac-Phe levels by using age- and sex-adjusted linear regression models to first relate Lac-Phe to metformin use (β = 0.88, P = 7.7×10^−10^) and then similar models further adjusted for baseline metformin use to relate Lac-Phe to BMI (β = −0.15, P = 6.4×10^−3^)(Fig. 4B). Consistent with a mediation effect of Lac-Phe, we observed that the effect of metformin use on BMI was weakened when Lac-Phe was included in the regression compared to the effect of metformin on BMI in the unmediated model (β = −0.39, P = 0.18) (Fig. 4B). The difference between the effect of metformin use on BMI in the unmediated and mediated models represents the mediation effect of Lac-Phe. As shown in Fig. 4C, using a bootstrapping method, we detected a significant mediation effect of Lac-Phe in the relationship between metformin use and BMI among MESA participants who lost weight during the study period (β = −0.13; 95% CI, −0.03 to −0.26; P = 5.2×10^−3^).

**Figure 4.**
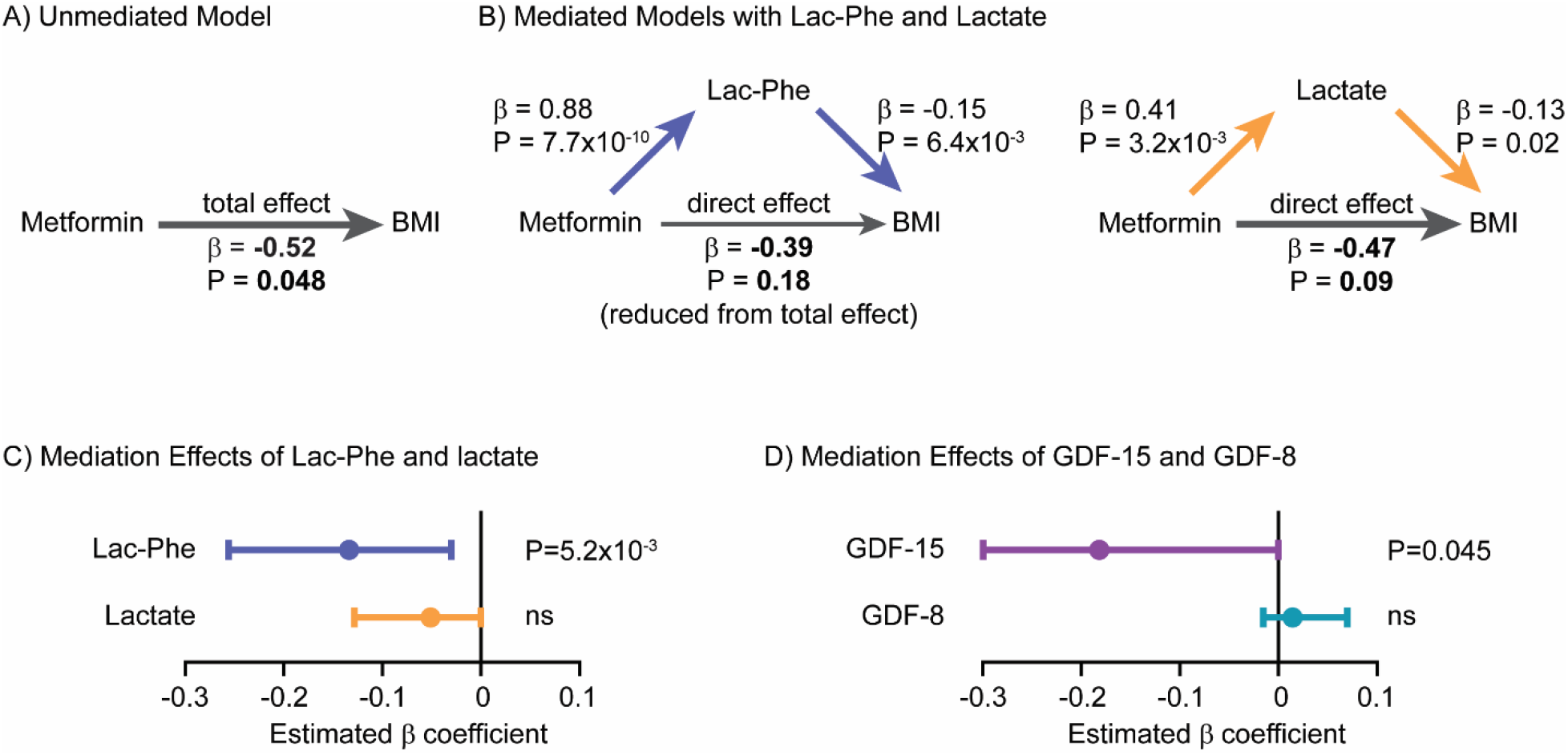
Lac-Phe mediates in part the effect of metformin on BMI reduction in a post-hoc subgroup analysis of MESA participants. (A) Among MESA participants who lost weight during the study period, the total effect of metformin use on BMI in was assessed in an “unmediated model” using an age- and sex-adjusted linear regression model. (B) To construct mediation models, the individual associations of metformin use, lac-phe, lactate, and BMI were assessed using linear regression models as described in Methods. The direct effect of metformin use on BMI was then assessed using an age- and sex-adjusted linear regression model adjusted for either lac-phe (left) or lactate (right). A reduction in the direct effect of metformin on BMI compared to the total effect of metformin on BMI in the unmediated model suggested partial meditation. (C) A nonparametric bootstrapping method was used to calculate the confidence intervals and statistical significance of the mediation effects of Lac-Phe and lactate. (D) The mediation effects of GDF-15 and GDF-8 on the effect of metformin use on BMI were also calculate using the same methods among MESA participants who lost weight during the study period.

Several control analyses demonstrated that the mediation effect of Lac-Phe on metformin-associated BMI reduction was specific. For example, we did not detect a significant mediation effect of lactate on metformin-associated BMI reduction (Fig. 4B, right). When we used the same set of pairwise associations but reordered the mediation model to test if BMI partially mediates the relationship between metformin use (exposure) and Lac-Phe levels (outcome), we obtained the expected null result (Fig. S4A-C). Among the subgroup of MESA participants who gained weight during the study period, we also did not detect a significant mediation effect of Lac-Phe or lactate on metformin-associated BMI increase (Fig. S4D-F). Finally, we expanded our analyses to test the mediation effects of each of the additional 136 metabolite levels that we recently measured in participants of MESA using the amide negative LC-MS method. As shown in Fig. S4G, we were unable to detect a significant mediation effect for the vast majority (129/136, 95%) of these metabolites.

Lastly, we applied similar mediation analysis in the subgroup of MESA participants who lost weight to determine the contribution of GDF15 to the effects of metformin on body weight. As shown in Fig. 4D, we also detected a partial mediation effect of GDF15 on metformin-associated BMI reduction. The mediation effect of GDF15 was comparable to that of Lac-Phe (Lac-Phe β = −0.13; 95% CI, −0.03 to −0.26; GDF15 β = −0.18; 95% CI, −0.00 to −0.41). In addition, the GDF15 mediation effect was also specific because we detected no significant mediation effect of the related hormone GDF8 (myostatin) on metformin-associated BMI reduction. Taken together, these post-hoc subgroup analyses support a causal role for Lac-Phe, as well as GDF15, in partially mediating the effects of metformin on body weight in humans.

### CNDP2-KO mice are resistant to the anti-obesity effects of metformin

To perform a direct and functional test of the contribution of Lac-Phe to the effects of metformin on body weight, we used global CNDP2-KO mice. These animals had previously been shown to have lower basal and exercise-inducible Lac-Phe levels and also exhibited increased food intake and body weight after a combined high fat diet/treadmill exercise training protocol^19^. CNDP2-KO mice were generated by heterozygotes breeding pairs and WT littermates were used as controls. Both WT and CNDP2-KO mice were given high fat diet for two months to induce obesity. As expected, a single dose of 300 mg/kg metformin treatment increased plasma Lac-Phe levels in WT mice. In CNDP2-KO mice, plasma Lac-Phe levels were dramatically reduced and not further increased following metformin treatment (Fig. 5A). Lactate levels remain the same after metformin treatment in both genotypes (Fig. S5A).

**Figure 5.**
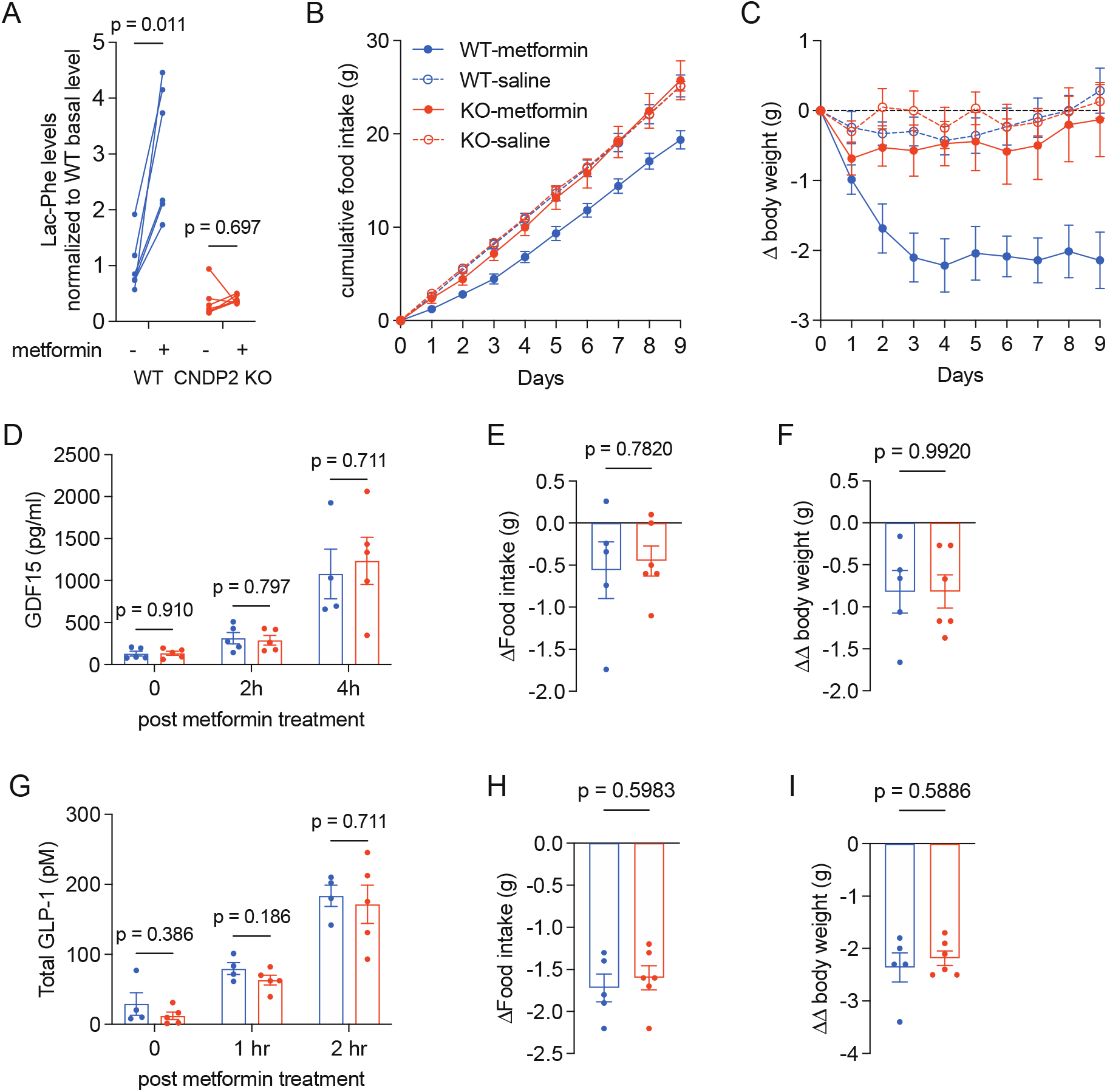
CNDP2 mediates the anti-obesity effects of metformin treatment in mice. (A) 300 mg/kg metformin treatment increases Lac-Phe levels in WT (n = 6), but not in CNDP2-KO (n = 7) mice. (B) Chronic metformin treatment (300 mg/kg daily) suppresses food intake of WT, but not CNDP2-KO mice. (C) Chronic metformin treatment (300 mg/kg daily) suppresses body weight of WT, but not CNDP2-KO mice. (D) 300 mg/kg metformin treatment increases circulating GDF15 levels in WT and CNDP2-KO mice. (E-F) 4 nmol subcutaneous GDF15 injection reduces food intake (E) and body weight (F) in WT and CNDP2-KO mice. Values adjusted by effects with vehicle treatment. (G) 300 mg/kg metformin treatment increases circulating GLP-1 levels in WT and CNDP2-KO mice. (H-I) 10 nmol subcutaneous Semaglutide injection reduces food intake (H) and body weight (I) in WT and CNDP2-KO mice. Values adjusted by effects with vehicle treatment. For (B) and (C), N = 6 for KO-saline, N = 7 for other groups. N = 4 – 5 in (D) and (G). N = 5 – 6 in (E), (F), (H), and (I). P values in (A) were calculated using multiple paired t tests with Holm-Šídák corrections; Welch t tests were used in (D-I).

Next, we treated a cohort of WT and CNDP2-KO mice with saline vehicle or with 300 mg/kg/day metformin (PO). At the beginning of this experiment, body weights were not different between genotypes (mean ± SEM: 38.4 ± 1.5 g for WT, 36.2 ± 1.6 g for CNDP2-KO mice, P = 0.32). Saline-treated WT and CNDP2-KO mice consumed 2.8 ± 0.1 g food per day and maintained their body weight throughout the experiment. As expected, metformin treatment of WT mice resulted in a durable suppression of food intake (mean ± SEM, 2.2 ± 0.1 g/day), resulting in a 2.1 ± 0.4 g (mean ± SEM) reduction in total body weight at the end of the experiment. By contrast, CNDP2-KO mice exhibited a food intake (mean ± SEM, 2.9 ± 0.2 g/day) and body weight change (mean ± SEM, −0.1 ± 0.5 g) that were not significantly different from saline-treated mice (P > 0.05). We conclude that CNDP2-KO mice are resistant to both metformin’s induction of Lac-Phe levels and also metformin’s effects on body weight and food intake.

To determine if Lac-Phe and CNDP2 also contributes to the anti-diabetic effect of metformin, we performed intraperitoneal glucose tolerance test (GTT) in WT and CNDP2-KO mice which had been on HFD for 8 weeks. Once again, basal body weights between genotypes at this point were not significantly different (mean ± SEM: WT, 41.1 ± 1.6 g; CNDP2-KO, 38.9 ± 1.5 g). After 4 hr fasting, all mice received a single dose of metformin (300 mg/kg, PO) simultaneously with glucose (1 g/kg, i.p) and blood glucose was measured over a 2 h period. WT and CNDP2-KO mice showed equivalent glucose levels at all time points (Fig. S5C,D). We conclude that the anti-diabetic effect of metformin is not affected in CNDP2-KO mice.

Metformin has been shown to induce the expression of multiple hormones which are involved in appetite control^9,10,12,18,26,27^. We measured the levels of GDF15, GLP-1, and PYY after metformin treatment in both WT and CNDP2-KO mice. 12-14 weeks old WT and CNDP2-KO mice were treated with one single dose of metformin (300 mg/kg, PO). In WT mice, plasma GDF15 levels increased from 130 ± 29 pg/ml (mean ± SEM, basal level) to 314 ± 68 pg/ml and 1078 ± 295 pg/ml at 2 and 4 hrs post treatment, respectively (Fig. 5D). An equivalent induction of GDF15 was observed in CNDP2-KO mice (Fig. 5D). Similarly, levels of GLP-1 and PYY were not different between WT and CNDP2-KO mice after metformin administration (Fig. 5G, S5B). Therefore the blunted effects of metformin on body weight in CNDP2-KO mice are not due to differences between genotypes in levels of these other feeding-regulating peptide hormones.

These data, together with our previous data^19^, demonstrate that CNDP2-KO mice are resistant to the anti-obesity effects of two distinct physiologic stimuli, exercise training and metformin treatment. To rule out the possibility that CNDP2-KO mice are more broadly resistant to any and all anorexigenic and anti-obesity stimuli, we treated WT and CNDP2-KO mice with recombinant GDF15 protein (4 nmol/kg, subcutaneous injection) or with the GLP-1 receptor agonist Semaglutide (10 nmol/kg, subcutaneous injection).^28^ In both experiments, CNDP2-KO mice showed comparable reduction in food intake (Fig. 5E,H) and body weight (Fig. 5F,I) as WT mice. These data demonstrate that CNDP2-KO mice have functional GDF15/GFRAL and GLP-1/GLP-1R pathways. In addition, they show that the resistance to anorexigenic stimuli in CNDP2-KO animals is limited to those perturbations that share a common feature of increased glycolytic flux and increased lactate mass action.

## Discussion

Here we show multiple lines of evidence demonstrating that Lac-Phe is a key downstream mediator of the effects of metformin on energy balance: 1) metformin is a strong pharmacological inducer of Lac-Phe levels in both mice and humans; 2) metformin increases Lac-Phe levels in a cell autonomous, CNDP2-dependent manner in macrophages and gut epithelial cells, but not in hepatocytes; and 3) global CNDP2-KO mice are resistant to metformin-inducible increases in Lac-Phe and also resistant to the anti-obesity effects of metformin therapy. These data designate the anorexigenic metabolite Lac-Phe as a key downstream mediator of the anti-obesity effects of metformin.

The precise relationship between GDF15, Lac-Phe, and metformin in energy balance remains an important area for future work. Our studies provide strong evidence from mice and humans that Lac-Phe is an important player in metformin’s effects on body weight. In addition, our statistical mediation analysis in humans would point to a role for both Lac-Phe and GDF15 as downstream mediators of metformin’s anti-obesity effects. The eventual identification of the molecular targets and neuronal circuits involved in the anorexigenic effects of Lac-Phe, and whether or not such pathways overlap with those defined by the GDF15/GFRAL pathway, would also help to clarify this question.

Our data, along with our previous study^19^, shows that CNDP2-KO mice are resistant to both the anti-obesity effects of exercise as well as that of metformin. However, CNDP2-KO mice are still fully responsive to other anorexigenic agents such as GLP-1 receptor agonists as well as to GDF15, two classes of molecules that themselves do not increase glycolytic flux or Lac-Phe levels. In addition, levels of GLP-1 and GDF15 are both elevated equivalently in WT and CNDP2-KO mice, establishing that the body weight phenotype of CNDP2-KO mice is not simply due to differences in the levels of these other feeding-associated hormones. Regardless, increased glycolytic flux, lactate metabolism, and Lac-Phe biosynthesis appears to be a hallmark of the types of anorexigenic stimuli where CNDP2-KO mice exhibit differences compared to WT mice in terms of food intake and body weight. Based on this conceptual framework, we predict that CNDP2-KO mice will also be resistant to the anti-obesity effects of sepsis and hypoxia therapy, but not lithium chloride. These are hypotheses that can be directly tested in future work.

We provide substantial data from cell culture studies that the biochemical mechanism by which metformin increases Lac-Phe involves inhibition of complex I and re-wiring of glycolytic flux to drive intracellular lactate mass action in CNDP2+ cells. Like sprint exercise, this biochemical mechanism once again relies on lactate mass action as the key energetic and kinetic determinant of Lac-Phe levels. However, in the case of metformin the origin of the lactate is from intracellular glycolytic pools in CNDP2+ cells, whereas in sprint exercise lactate is coming from extracellular, muscle-derived sources. This intracellular versus extracellular difference in origin of lactate also therefore explains why in sprint exercise elevates both lactate and Lac-Phe levels, whereas with metformin treatment only increases Lac-Phe levels, without causing concomitant increases in circulating lactate as well.

Cells expressing the Lac-Phe biosynthetic enzyme CNDP2 include macrophages, other immune cells, as well as epithelial cells of the kidney, gut, and lung. We suspect that gut epithelial cells and intestinal macrophages are likely primary cell types responsible for Lac-Phe biosynthesis in response to metformin. This is because the concentrations of metformin needed to inhibit complex I are not reached systemically, but clearly are sufficiently high in the local environment of the gut. Such a model would also suggest that gut-resident macrophages can exert endocrine functions, which is an interesting idea that warrants additional studies. Whether or not these are the same CNDP2+ cell populations responsible for Lac-Phe biosynthesis in response to exercise remains an unanswered question. In the context of exercise, where systemic levels of lactate are elevated, other non-gut CNDP2+ cell types may be larger contributors to the pool of circulating Lac-Phe. Conditional *Cndp2* alleles, which are actively being developed in our laboratory, should enable functional dissection of the relative contributions of specific CNDP2+ cell types to Lac-Phe biosynthesis in response to these diverse stimuli.

## Methods

### Chemicals

Metformin hydrochloride (13118), buformin hydrochloride (18507), phenformin (14997), and Semaglutide acetate (29969) were purchased from Cayman Chemical (13118). Proguanil hydrochloride (PHR1713), D-Glucose-^13^C_6_ (389374), 2,4-Dinitrophenol (D198501), Rotenone (R8875), 2-Thenoyltrifluoroacetone (T27006), and Sodium azide (S2002) were purchased from Sigma-Aldrich. Oligomycin (AAJ61898MA) was purchased from Fisher Scientific. Recombinant GDF15 (957-GD) was purchased from R&D systems.

### Cell culture

Cell lines used in this study were obtained from the American Type Culture Collection (ATCC) and grown at 37 °C with 5% CO_2_. BV-2 cells were a gift from Dr. Wyss-Coray. Caco-2, BV-2, C2C12, 3T3-L1, SW48, HEK293T, and AML12 cells were grown in DMEM (Corning 10-017-CV) with 10% FBS (Corning 35-010-CV) and 1% Penicillin-Streptomycin (Pen-Strep, Gibco 15140). AML12 cells were also supplemented with Insulin-Transferrin-Selenium (ITS-G, Gibco 41400). Neuro2A and HepG2 cells were grown in EMEM (Corning 10-009-CV) with 10% FBS and 1% Pen-Strep.

Primary peritoneal macrophages were collected as described in Zhang et al.^29^,with slight modifications. Briefly, peritoneal macrophages were harvested with ice-cold PBS 3-5 days post *i.p.* injection of 1 mL of 3% Brewer thioglycolate medium. The cells were later plated in DMEM/F-12 (Gibco 11320) medium supplemented with 10% FBS, 1% Pen-Strep, and 10 mM L-Glutamine (ATCC, 30-2214). Primary hepatocytes were isolated following perfusion, digestion, and Percoll gradient centrifugation as in Jung et al.^30^.

### Generation of CNDP2-KO cell lines

CNDP2-KO Caco-2 and BV-2 cells are generated using the pLentiCRISPRv2 system^31^. For the mouse cell lines, the single guide RNA (sgRNA) used was 5′-CAGTGAAATGAGATCCGTCA-3′. For human cell lines, the sgRNA used was 5′-ACAGAAACTCGCAAAATGGG-3′. As per the protocols described by the Zhang group, oligonucleotides for the sgRNA and reverse complement sequences were synthesized and cloned into the plentiCRISPRv2 vector. For the mouse cell lines, the following oligonucleotides were used: fwd, 5′-CACCGCAGTGAAATGAGATCCGTCA-3′; rev, 5′-AAACTGACGGATCTCATTTCACTGC-3′. For the human cell lines, the following oligonucleotides were used: fwd, 5′-CACCGACAGAAACTCGCAAAATGGG-3′; rev, 5′-AAACCCCATTTTGCGAGTTTCTGTC-3′. Lentivirus particles were generated in the HEK293T cell line using Polyfect for the co-transfection of the cloned plentiCRISPRv2 plasmid with the viral packing psPAX2 plasmid and the viral envelope pMD2.G plasmid. A parental plentiCRISPRv2 plasmid was used as a control. Lentiviral supernatants were collected after 24 h and filtered through a 0.45-µM filter. The supernatant was then mixed in a 1:1 ratio with polybrene to a final concentration of 8 µg/ml polybrene. This mixture was added to cells at 40–50% confluence in 6-well plates. Transduced cells were transferred to a 10-cm plate followed by selection by puromycin for 1 week. Caco-2 and BV-2 cells were selected with 10 ug/ml and 5 ug/ml puromycin, respectively. Knockout of CNDP2 was verified by Western Blot. Antibodies used were described below: rabbit anti-CNDP2 (Proteintech, 14925-1-AP), rabbit anti-beta tubulin (Origene, TA301569), goat anti-rabbit IgG (LI-COR Biosciences, 926-32211).

### Lac-Phe *in vitro* production

For cell lines, cells were plated in 12-well plates at 70–80% confluence. The next day, cells were washed two times with PBS and incubated in 0.5 ml medium supplemented with various concentrations of metformin as indicated. After overnight incubation, 400 μl of medium was transferred to a new Eppendorf tube and 20 μl 1 M hydrochloric acid was added to acidify the medium and to protonate Lac-Phe. 1:1 v: v ethyl acetate was added into each sample and vortexed for 30 s to extract Lac-Phe into the organic layer. A total of 300 μl from the top layer was transferred to a new Eppendorf tube and dried down under a stream of nitrogen. The residue was re-suspended in 100 µl of a 2:1:1 mixture of acetonitrile: methanol: water. Cells were kept on ice to collect the lysate. Cells were disassociated with trypsin and collected into an Eppendorf tube. 200 ul PBS was used to wash each well to ensure all cells were collected. Cells were then centrifuged at 4 °C for 10 min at 5,000 r.p.m. and the supernatant removed to obtain the cell pellet. A volume of 100 μl of a 2:1:1 mixture of acetonitrile: methanol: water mixture was used to lyse the cells and precipitate large proteins. The mixture was centrifuged at 4 °C for 10 min at 15,000 r.p.m. and the supernatant was transferred to a LC–MS vial.

### Cellular respiration measurements

Cellular oxygen consumption rates were determined using Agilent XF96 Analyzer assay was performed following Agilent Seahorse XF Cell Mito Stress Test Kit manual, with slight modifications. Briefly, cells were plated at 100k per well in the XF96 cell culture plate, and incubated with metformin at various concentrations in growth media overnight. The next morning, the cells were washed with PBS and incubated with Seahorse assay buffer (8.3 g/L DMEM (Sigma D5030), 1.8g/L NaCl, 1 mM pyruvate, 20 mM glucose, 1% Pen-Strep, pH 7.4). Final concentrations of compounds were used as follows: oligomycin, 10 μM; DNP, 100 μM; rotenone, 3 μM.

### Lactate and Lac-Phe tracing using Glucose-^13^C_6_

^13^C glucose or ^12^C glucose of the same molar concentration was added to Glucose-free DMEM (Gibco A1443001) supplemented with 10% FBS, 1% Pen-Strep, 10 mM L-glutamine. Cells plated in 12-well plate was washed with PBS twice and cultured in ^13^C or ^12^C glucose media overnight. Lactate and Lac-Phe with various numbers of ^13^C isotopes were detected with the following *m/z*: 0: 89.0244, 236.0928; 1: 90.0278, 237.0962; 2: 91.0311, 238.0995; 3: 92.0345, 239.0129; 4: 93.0378, 240.1062.

### Lac-Phe measurement adding ^13^C_3_-lactate

Primary mouse macrophages were plated in 12-well plate at 1 million per well. 10 mM ^13^C-labeled sodium lactate was added to the cells with or without 5 mM metformin overnight. Lac-Phe was detected with LC-MS.

### General animal Information

All animal experiments were performed according to procedures approved by the Stanford University Administrative Panel on Laboratory Animal Care. Mice were maintained in 12-hour light-dark cycles at 22 °C and 50% relative humidity and fed a standard irradiated rodent chow diet. Where indicated, a high-fat diet (60% kcal from fat, Research Diets, D12492) was used. C57BL/6J (000664) and C57BL/6J DIO (380050) mice were purchased from the Jackson Laboratory. CNDP2-KO mice were as previously described^32^, and originally obtained from the Mutant Mouse Regional Resource Center, a NCRR-NIH funded repository. CNDP2-KO mice used in this study were generated by heterozygous breeding crosses, as in [Li et al., 2023]. The following genotyping primers were used: WT allele (fwd, 5′-CAGATGGCTCGGAGATACCAC-3′; rev, 5′-TTCCCGCTCCACCAAGGTGAAG-3′); KO allele (fwd, 5′-GCTCTGTAAGGGAAAGAGATGACCC-3′; rev, 5′-AATAGGACATACCCAGTTCTGTGAGG-3′).

### Mouse plasma sample preparation for LC-MS analysis

Blood was collected from mice through a submandibular bleeding into lithium heparin tubes (BD, 365985) and immediately kept on ice. The blood was then centrifuged at 4 °C for 5 min at 5,000 r.p.m. Plasma was transferred into new Eppendorf tubes and frozen at −80 °C if not used immediately. Metabolites for LC-MS analysis were extracted by adding 150 ul of 2: 1 mixture of acetonitrile: methanol to 50 ul plasma. The mixture was then mixed by vortex and centrifugated at 4 °C for 10 min at 15,000 r.p.m. The supernatant was transferred to a LC-MS vial.

### Study Populations

The Stanford cohort of participants in Studies of Insulin Resistance and Diabetes was described in details previously^21^. The original study was conducted from 1998 to 2004. The study was approved by Stanford Institutional Review Board, and each individual gave written informed consent to participate in the study. Of the 31 participants in the original study, 21 had plasma samples available and were included in the current study. All participants had type 2 diabetes with fasting plasma glucose concentration greater than 170 mg/dL and were being treated with medical nutrition therapy alone (N = 10) or with sulfonylurea monotherapy (glipizide or glyburide 10 to 20 mg daily; N = 11). The dose of metformin started from 500 mg twice daily with slow uptitration to a final dose of 2 g daily. All patients received metformin treatment for 12 weeks. Baseline and post-treatment samples were collected after an overnight fast. The metformin treatment group consisted of 8 females and 13 males. Of those, 13 participants were white, 4 East Asian, 2 black, and 2 South Asian. At baseline, participants had mean ± SD age of 58 ± 10 years; BMI, 29.4 ± 4.6 kg/m2; weight 87.7 ± 25.1 kg; and fasting glucose, 234 ± 39 mg/dL. After 12 weeks of treatment with metformin, fasting glucose decreased to 182 ± 48 mg/dL (P <0.001), but weight remained unchanged 87.7 ± 28.5 kg (P = 0.88).

MESA is a U.S. community-based cohort study that recruited 6814 individuals who self-identified as White, African American, Hispanic, or Chinese American, as previously described^22^. Included in the present study are 3656 individuals across all four racial/ethnic groups with metabolomic profiling at the baseline examination (2000–2002). Baseline characteristics are described in Table S1. 3645 individuals had available follow-up BMI values from exam 5 (2010-2012) with a mean follow-up of 9.46 years. The MESA human protocol was approved by The Lundquist Institute (formerly Los Angeles BioMedical Research Institute) at Harbor-University of California, Los Angeles Medical Center, University of Washington, Wake Forest School of Medicine, Northwestern University, University of Minnesota, Columbia University, Johns Hopkins University, and University of California, Los Angeles Institutional Review Boards, and all participants provided written informed consent.

### Metabolite measurements by LC-MS

In murine studies and the Stanford cohort human studies, Plasma samples were collected and kept at −80 °C until preparation for LC-MS. 150 ul of 2: 1 mixture of acetonitrile: methanol was added to 50 ul plasma. The mixture was then mixed by vortex and centrifugated at 4 °C for 10 min at 15,000 r.p.m. The supernatant was transferred to a LC-MS vial. Untargeted metabolomics measurements were performed using an Agilent 6520 or 6545 Quadrupole time-of-flight LC–MS instrument. For lactate and Lac-Phe, MS analysis was performed using electrospray ionization (ESI) in negative mode. The dual ESI source parameters were set as follows: the gas temperature was set at 250 °C with a drying gas flow of 12 l/min and the nebulizer pressure at 20 psi; the capillary voltage was set to 3,500 V; and the fragmentor voltage set to 100 V. Separation of metabolites was conducted using a Luna 5 μm NH2 100 Å LC column (Phenomenex 00B-4378-E0) with normal phase chromatography. Mobile phases were as follows: buffer A, 95:5 water:acetonitrile with 0.2% ammonium hydroxide and 10 mM ammonium acetate; buffer B, acetonitrile. The LC gradient started at 100% B with a flow rate of 0.2 ml/min from 0 to 2 min. The gradient was then linearly increased to 50% A and 50% B at a flow rate of 0.7 ml/min from 2 to 20 min. From 20 to 25 min, the gradient was maintained at 50% A and 50% B at a flow rate of 0.7 ml/min.

For Metformin, MS analysis was performed using ESI in positive mode. The dual ESI source parameters were set as follows: the gas temperature was set at 325 °C with a drying gas flow of 5 l/min and the nebulizer pressure at 30 psi; the capillary voltage was set to 3,500 V; and the fragmentor voltage set to 175 V. Separation of metabolites was conducted using a Luna 5 μm C5 100 Å LC column (Phenomenex 00B-4043-E0) with reverse phase chromatography. Mobile phases were as follows: buffer A, water with 0.1% formic acid; buffer B, acetonitrile with 0.1% formic acid. The LC gradient started at 95%A and 5% B with a flow rate of 0.2 ml/min from 0 to 2 min. The gradient was then linearly decreased to 5% A and 95% B at a flow rate of 0.7 ml/min from 2 to 7 min. From 7 to 12 min, the gradient was maintained at 5% A and 95% B at a flow rate of 0.7 ml/min.

Quantification of the metabolite concentrations were performed by generating a standard curve with known concentrations of each metabolite. Metabolite standards were analyzed alongside the samples using the same method. A standard curve generated from the metabolite concentrations and extracted ion intensities were used to calculate the concentrations of each metabolite.

In MESA, metabolomics profiling was performed using LC-MS on fasting baseline plasma samples, as previously described^33,34^. Briefly, metformin and other water-soluble, polar metabolites were measured using a Nexera X2 U-HPLC (Shimadzu) equipped with a 150 x 2 mm, 3 μm Atlantis hydrophilic interaction LC column (Waters) coupled to a Q Exactive hybrid quadrupole Orbitrap MS (ThermoFisher Scientific). Metabolites were extracted from 10 μl plasma by adding 90 μl of Acetonitrile:Methanol:Formic acid (74.9:24.9:0.2,v/v/v) solution spiked with valine-d8 (Sigma) and Phenylalanine-d8 (Cambridge Isotope Laboratories). The metabolites were eluted at 0.25 ml/min with 5% buffer A (10 mM Ammonium-Formate, 0.1% formic acid in water) for 0.5 minutes followed by a linear gradient to 40% buffer B (0.1% formic acid in acetonitrile) over 10 minutes. MS analyses were carried out using electrospray ionization in the positive mode and full scan spectra were acquired over 70-800 m/z. Raw data were processed using Trace Finder (v3.3, Thermo Fisher Scientific) and Progenesis QI (Waters). Lac-phe, lactate, and other intermediary metabolites were measured using a 1290 Infinity LC system (Agilent Technologies) equipped with a 100 x 2.1 mm XBridge amide column (Waters) coupled to a 6490 Triple Quad MS (Agilent Technologies) in negative ionization mode via multiple reaction monitoring (MRM) scanning. Data were quantified using MassHunter Quantitative Analysis software (V10.1, Agilent).

To ensure quality control, a mixture of ∼150 reference standards was analyzed before, during periodic intervals throughout, and after each MS run to ensure reproducibility of LC retention times, LC peak shapes, and MS sensitivity. Isotope labeled internal standards were monitored in each sample throughout the duration of each run. Pooled plasma samples were monitored after every 10 participant samples to standardize for MS drift over time using “nearest neighbor” normalization and between batches. Separate pooled plasma samples were monitored after every 20 participant samples to determine coefficient of variation (CV) for each metabolite. Metabolite identities were confirmed using authentic reference standards. All metabolite peaks were manually reviewed for peak quality in a blinded manner. None of the included metabolites had poor peak quality or CVs ≥ 30% averaged across batches

### GDF-15 and GDF-8 Measurement in MESA

In MESA, proteomic profiling was performed using the Olink Explore platform, as previously described^35^. Relative quantification of GDF-15 (OID20251) and GDF-8 (OID20115) using the Olink Explore 3072 platform were available in 907 participants from the baseline exam that were randomly selected across all four racial/ethnic groups and that had available metformin LC-MS values from the baseline exam and BMI values from the baseline exam and exam 5.

### Animal experiments

#### Effect of various doses of metformin on food intake and body weight

C57BL/6J DIO mice on HFD were single housed 3 - 5 days prior to experiments and were mock *p.o.* injected with saline daily till body weights of all mice stabilized, to prevent stress-induced effects on food intake and body weight. The mice were then divided into groups of 5 – 6, and were *p.o.* treated with saline, or different concentrations of metformin hydrochloride as indicated in the figures for a period of 7 days. Metformin hydrochloride was directly dissolved in saline. Food intake and body weight were taken daily.

#### Effect of 300mg/kg metformin treatment on food intake and body weight in WT and CNDP2-KO mice

WT and CNDP2-KO mice were generated by heterozygous breeding crosses. At 4 – 6 weeks, mice were switched from chow diet to HFD. At 10 – 14 weeks, mice were single housed and mock *p.o.* injected with saline 3 – 5 days till body weight stabilized to reduced stress-induced loss of food intake and body weight. WT and CNDP2-KO were body weight matched and received either saline or 300 mg/kg metformin hydrochloride. Body weight and food intake were measured daily for a period of 9 days.

#### Effect of 4 nmol/kg GDF15 treatment on food intake and body weight in WT and CNDP2-KO mice

12 – 16 weeks old DIO WT and CNDP2-KO mice were single housed and mock *s.c* injected with vehicle (8: 1: 1 saline: DMSO: Kolliphor) for two days. On the experiment day, mice receive either vehicle or 4 nmol/kg GDF15 subcutaneously. Food intake and body weight were measured before and 24 hrs after the treatment.

#### Effect of 10 nmol/kg semaglutide treatment on food intake and body weight in WT and CNDP2-KO mice

12 – 16 weeks old DIO WT and CNDP2-KO mice, used in GDF15 study were allowed to recover for 3 days. The mice were then mock *s.c.* injected once. On the experiment day, mice receive either vehicle or 10 nmol/kg semaglutide subcutaneously. Food intake and body weight were measured before and 24 hrs after the treatment.

#### Hormone measurements after metformin treatment

12 – 14 weeks old WT and CNDP2-KO mice were treated with one single dose of 300 mg/kg metformin hydrochloride and blood were taken at time points indicated in the corresponding figures. Levels of hormones were measured with commercial ELISA kits following the manufacturers’ instructions. GLP-1 was measured after 4 hours fasting. Information about the ELISA kits are provided below: mouse PYY ELISA kit (81501) from Crystal Chem; mouse GLP-1 ELISA kit (EZGLP1T) from Sigma-Aldrich; mouse GDF15 ELISA kit (DY6385) from R&D systems.

#### Glucose tolerance test (GTT) in WT and CNDP2-KO mice

12 – 14 weeks old WT and CNDP2-KO mice were single housed and fasted for 4 hours. The mice then injected *i.p.* 1g/kg glucose and *p.o.* 300 mg/kg metformin hydrochloride. Blood glucose was measured at 0, 20, 40, 60, 90, 120 minutes post the treatment.

### Statistical Analyses

In MESA, metabolite levels were log transformed and standardized to have a mean of 0 and a SD of 1 within each batch. Metformin use was assessed by self-reported use of an oral hypoglycemic agent by study participants and validated by detection of plasma metformin levels by LC-MS during the baseline exam (AU > 50). Mediation analyses were performed using the framework originally suggested by Baron and Kenny^36^ and others^37,38^. Briefly, a sequence of linear regressions were performed to test if Lac-Phe or lactate levels were predicted to partially mediate the effect of metformin use (exposure) on BMI (outcome) in each MESA subgroup (participants who lost weight and participants who gained weight during the study period). Participants who had available metformin, Lac-Phe, and lactate levels measured by LC-MS, as well as BMI values at Exam 1 and Exam 5 were included in the analyses. In step one of the analysis, the association between metformin use and BMI was assessed using age- and sex-adjusted linear regression models. This association formed the basis for the total effect between metformin use and BMI in an “unmediated model”. In step 2, Lac-Phe or lactate levels were related to metformin use using an age- and sex-adjusted linear regression model. In step 3, Lac-Phe or lactate levels were related to the difference in BMI between Exams 1 and 5 using an age- and sex-adjusted linear regression model that was further adjusted for baseline metformin use. This additional adjustment for metformin use was included in order to identify the association between Lac-Phe (or lactate) and BMI that was independent of the shared correlation between each variable and the exposure of metformin use. In order to perform a mediation analysis, each association between exposure and outcome, exposure and mediator, mediator and outcome was required to be significant with p-value ≤ 0.05. Mediation analyses were conducted by regressing BMI (outcome) on metformin use (exposure) with adjustment for the mediator (e.g., Lac-Phe or lactate levels). The difference between the effect of metformin use on BMI in the unmediated and mediated models represented the mediation effect. The 95% confidence interval and statistical significance of the mediation effect was calculated using a nonparametric bootstrapping method, as previously described^39,40^ using the R package mediate^41^.

## Acknowledgements

We thank members of the Long lab for helpful discussions. We thank Dr. Wyss-Coray for sharing the BV-2 microglial cell line. We thank Lichao Liu from the Wu lab for helping with Seahorse assays. Dr. Benson acknowledges supported by NHLBI K08HL145095. This study was also supported in part by the National Heart, Lung and Blood Institute (NHLBI) TOPMed MESA Multi-Omics (HHSN2682015000031/HSN26800004). The MESA projects are conducted and supported by the National Heart, Lung, and Blood Institute (NHLBI) in collaboration with MESA investigators. Support for the Multi-Ethnic Study of Atherosclerosis (MESA) projects are conducted and supported by the National Heart, Lung, and Blood Institute (NHLBI) in collaboration with MESA investigators. Support for MESA is provided by contracts 75N92020D00001, HHSN268201500003I, N01-HC-95159, 75N92020D00005, N01-HC-95160, 75N92020D00002, N01-HC-95161, 75N92020D00003, N01-HC-95162, 75N92020D00006, N01-HC-95163, 75N92020D00004, N01-HC-95164, 75N92020D00007, N01-HC-95165, N01-HC-95166, N01-HC-95167, N01-HC-95168, N01-HC-95169, UL1-TR-000040, UL1-TR-001079, UL1-TR-001420, UL1TR001881, DK063491, and R01HL105756. The authors thank the other investigators, the staff, and the participants of the MESA study for their valuable contributions. A full list of participating MESA investigators and institutes can be found at http://www.mesa-nhlbi.org. Supported in part by the National Institutes of Health, National Heart, Lung, Long and Blood Institute (NHLBI) contract 1R01HL151855 and the National Institute of Diabetes and Digestive and Kidney Diseases contract UM1DK078616.

## Supplementary figures and figure legends

**Supplementary Figure 1.**
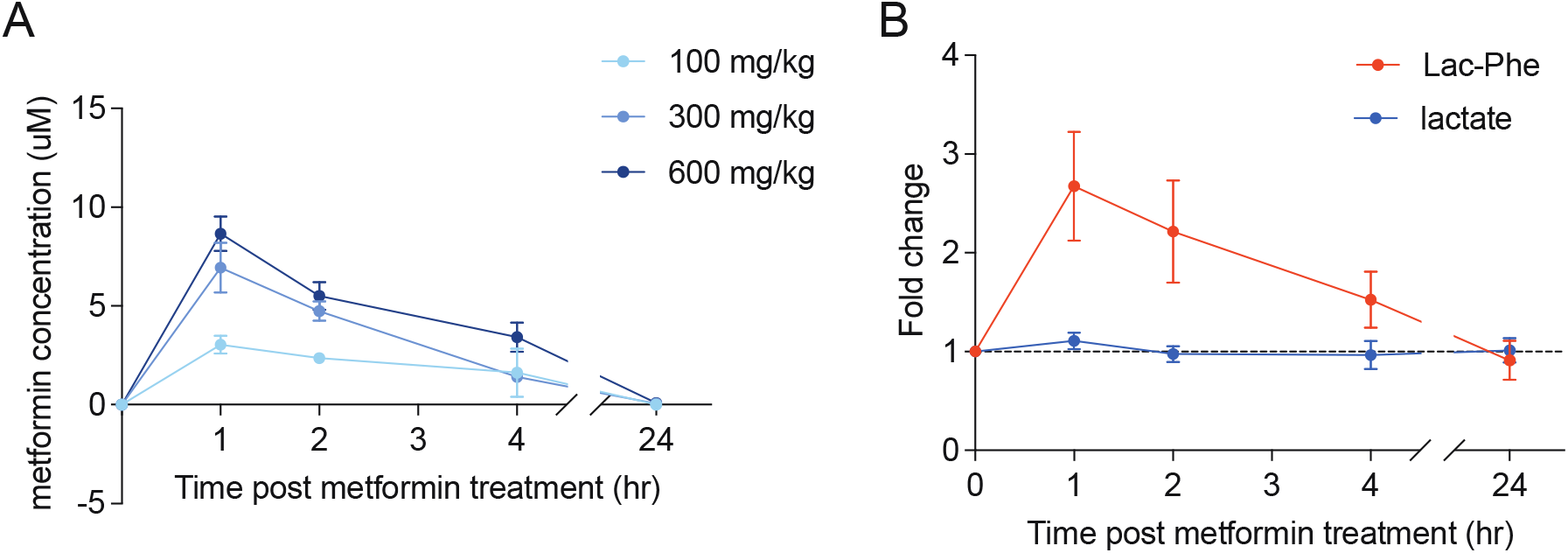
Plasma metabolite concentrations after a single dose of metformin in mice. (A) Metformin levels in the plasma after the mice were gavaged with 100, 300, or 600 mg/kg metformin (n = 5). (B) Fold changes of lactate and Lac-Phe after 300 mg/kg metformin (P.O) (n = 5).

**Supplementary Figure 2.**
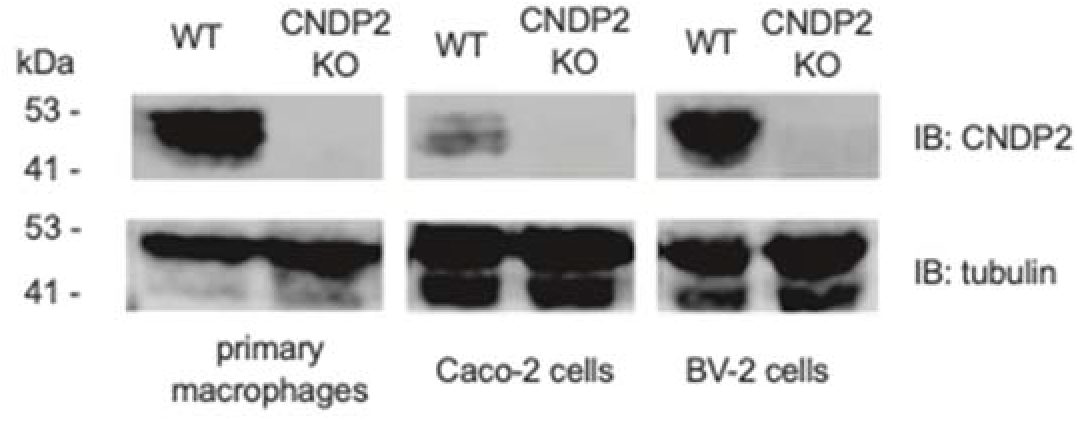
Western blot of CNDP2-KO and control cells. Western blot was performed using primary antibodies against CNDP2 and beta-tubulin. The experiment was performed twice, similar results were obtained.

**Supplementary Figure 3.**
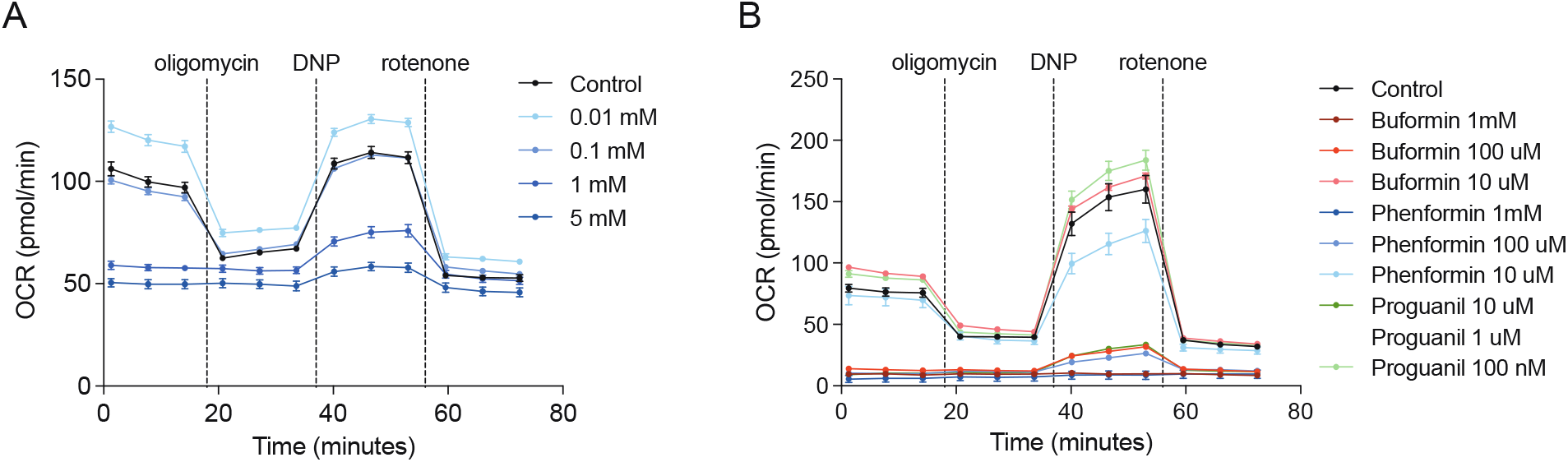
Metformin and other biguanides inhibit cellular mitochondrial respiration. (A) Metformin inhibits mitochondrial respiration in Caco-2 cells. (B) Other biguanides inhibit mitochondrial respiration in primary macrophages. N = 5 per group.

**Supplemental Figure 4.**
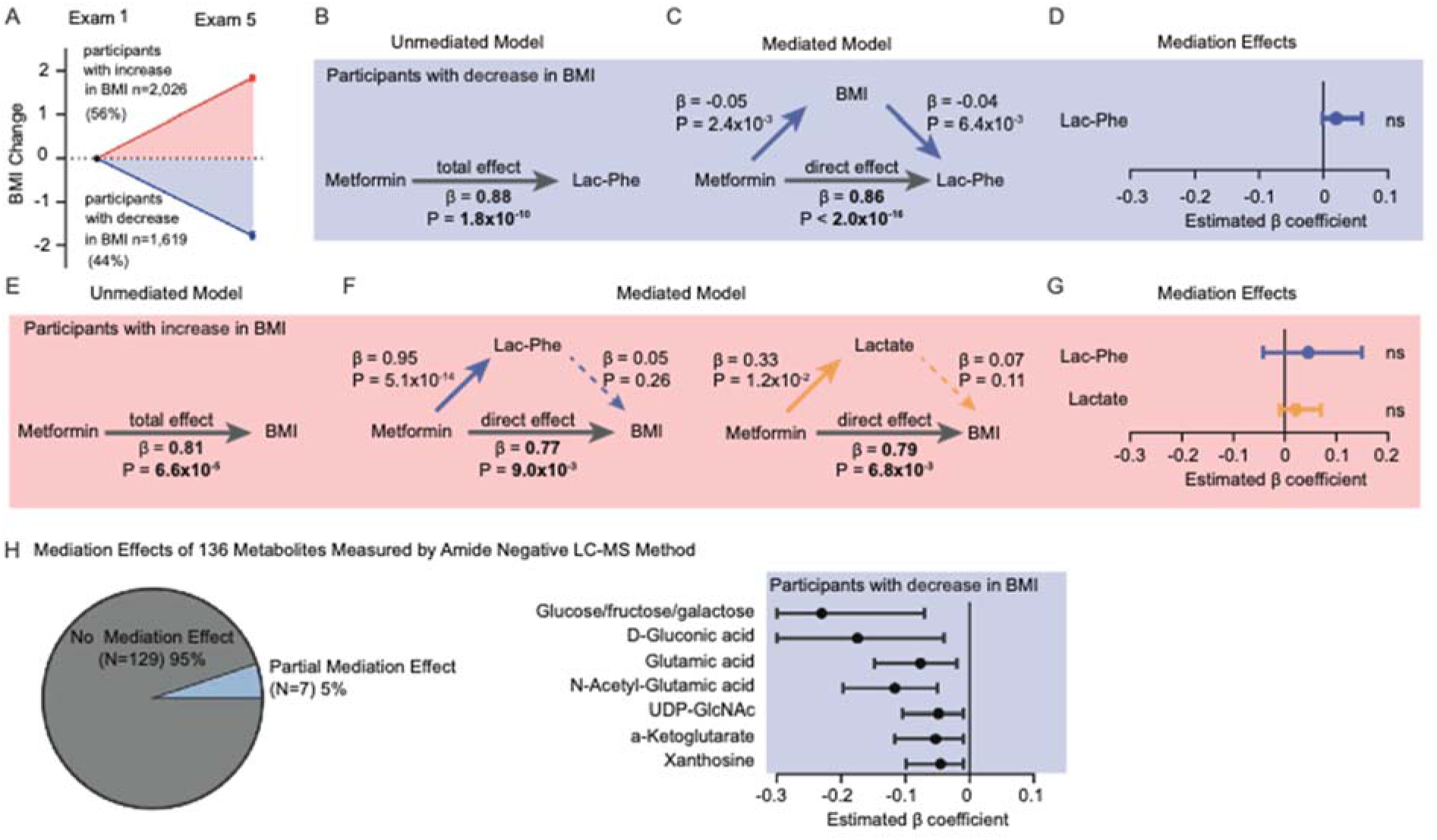
Control analyses for the mediation effect of Lac-Phe on metformin-associated BMI reduction. (A) Two subgroups in the post-hoc subgroup analysis. The change in BMI of the complete MESA sample prior to subgroup analysis was 0.24 ± 0.04 kg/m^2^, n = 3645. Participants with increased BMI are shown in red; participants with decreased BMI are shown in blue. (B-D) The mediation model in Fig.4B was reordered to test if BMI partially mediates the relationship between metformin use (exposure) and Lac-Phe levels (outcome). (B) Among MESA participants who lost weight during the study period, the total effect of metformin use on Lac-Phe levels was assessed in an “unmediated model” using an age- and sex-adjusted linear regression model. (C) To construct mediation models, the individual associations of metformin use, lac-phe, lactate, and BMI were assessed using linear regression models as described in Methods. The direct effect of metformin use on Lac-Phe levels was then assessed using an age- and sex-adjusted linear regression model adjusted for BMI. No reduction in the direct effect of metformin on Lac-Phe levels was appreciated compared to the total effect of metformin on Lac-Phe in the unmediated model suggesting no meditation. (D) A nonparametric bootstrapping method was used to calculate the confidence intervals and statistical significance of the mediation effects of BMI. Next, the mediation effects of Lac-Phe and lactate on the effect of metformin-associated BMI increase was assessed (E-G). (E) Among MESA participants who gained weight during the study period, the total effect of metformin use on BMI was assessed in an “unmediated model” using an age- and sex-adjusted linear regression model. (F) To construct mediation models, the individual associations of metformin use, lac-phe, lactate, and BMI were assessed using linear regression models as described in Methods. The direct effect of metformin use on BMI was then assessed using an age- and sex-adjusted linear regression model adjusted for either lac-phe (left) or lactate (right). No reduction in the direct effects of metformin on BMI compared to the total effect of metformin on BMI in the unmediated model suggested no meditation effect of either Lac-Phe or lactate. (G) A nonparametric bootstrapping method was used to calculate the confidence intervals and statistical significance of the mediation effects of Lac-Phe and lactate. (H) Finally, the mediation effect of each of the additional 136 metabolite levels measured in participants of MESA using the amide negative LC-MS method was assessed on metformin-associated BMI reduction among participants who lost weight during the study period. Seven out of the 136 tested metabolites were found to have a predicted mediation effect with unadjusted p-value ≤ 0.05. The median values and 95% confidence intervals of the predicted mediation effects for these seven metabolites were calculated using a nonparametric bootstrapping method, as above.

**Supplementary Figure 5.**
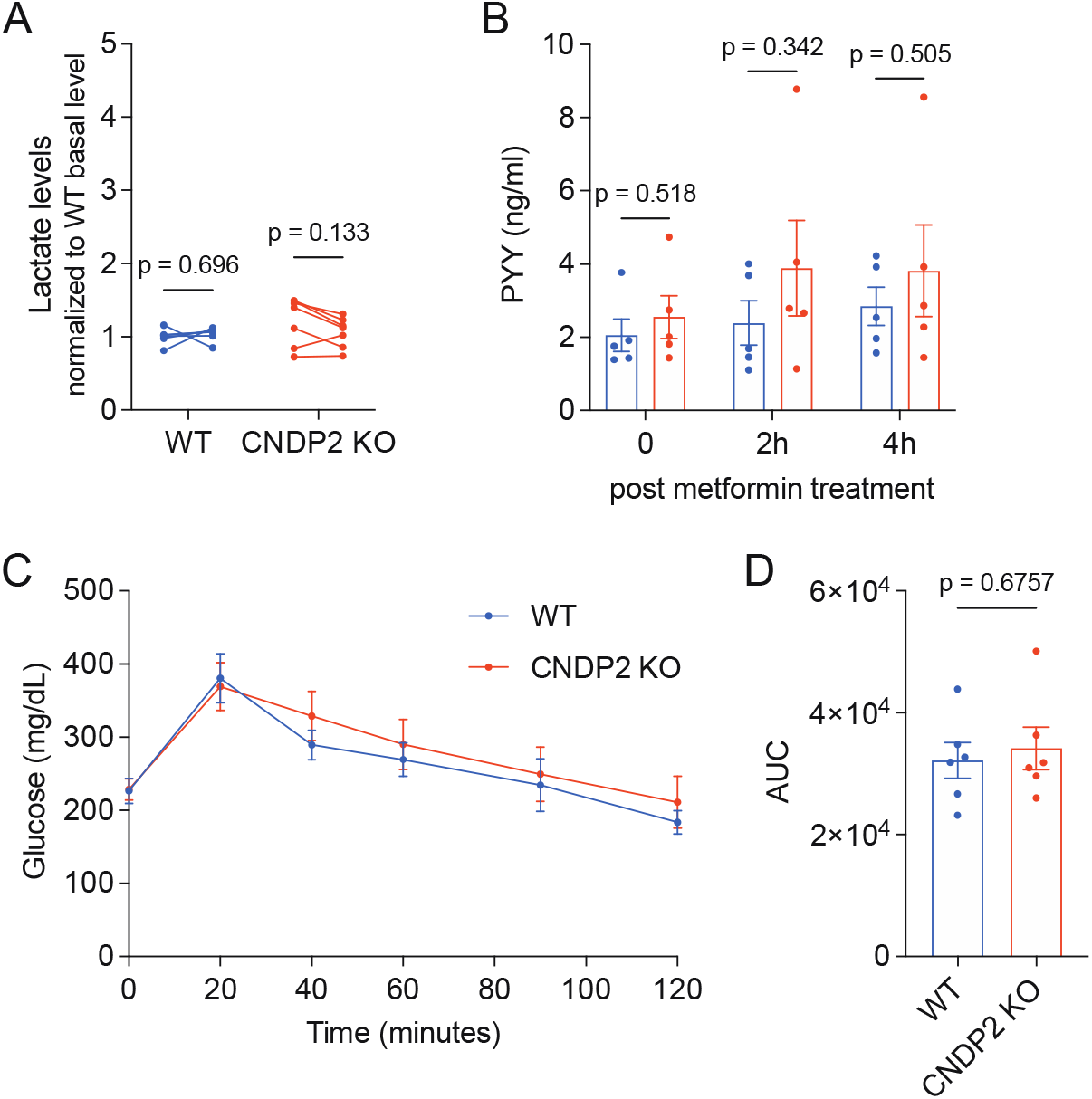
CNDP2 mediates anti-obesity effects, but not anti-diabetic effects of metformin. (A) Lactate levels in WT and CNDP2-KO mice after a single dose of 300 mg/kg metformin treatment. (B) Circulating PYY levels in WT and CNDP2-KO mice after 300 mg/kg metformin treatment. (C) Glucose levels of WT and CNDP2-KO mice during GTT after receiving 300 mg/kg metformin. (D) Area under curve plot of results shown in (C). For (A), N = 6 for WT, N = 7 for CNDP2-KO. Multiple paired t tests with Holm-Šídák corrections; In (B), N = 5, Welch t tests were used. N = 6 in (C-D). Two-way RM ANOVA was used in (C), Welch t test was used in (D).

